# Human umbilical mesenchymal stem cell-derived mitochondria ameliorate maternal phenotype by improving placental mitochondria and vascular function in angiotensin II-induced preeclampsia rat

**DOI:** 10.1101/2024.05.08.592162

**Authors:** Hui Xing Cui, Jun Xian Liu, Young Cheol Kang, Kyuboem Han, Hong Kyu Lee, Chun-Hyung Kim, Yin Hua Zhang

## Abstract

**Background:** Mitochondrial transplantation (Mito-T) is a novel therapeutic strategy targeting ischemic cardiovascular diseases. Here, we tested the efficacy of human umbilical mesenchymal stem cell-derived mitochondria transplantation (Mito-T) on a rat model of PE.

**Methods:** PE was induced by infusing angiotensin II (Ang II) to SD pregnancy rats on gestation day 8 (GD 8). Mito-T (100 μg/μl) was injected *via* jugular vein on GD 14.

**Findings:** On GD 20, PE rats showed high blood pressure, kidney and placental vascular abnormalities, reduced placental and fetal weights. Injected Mito-T was distributed intensively in the kidney, uterus and placenta of PE rats. Importantly, Mito-T reversed clinical manifestations of PE, restored placental abnormalities and reduced serum sFLT-1 levels and sFLT-1/PlGF ratio. In the placental mitochondria, Mito-T increased ETC complexes (complex I-V), improved mitochondrial membrane potential, ATP synthase and citrate synthase activities and biogenesis markers (PGC-1α, TFAM, NRF1) and reduced ROS production. Mito-T increased mitochondrial fusion proteins (OPA1, MFN1 and MFN2), reduced mitochondrial fission proteins (DRP1 and FIS1) and mitophagy proteins (PINK, BNIP3, BNIP3L, FUNDC1), restored sFLT-1 regulating calcineurin-NFAT-dependent pathways in the placental tissue, primary trophoblast cells and Bewo cell line. Furthermore, eNOS, nNOS and AT2R mRNA and protein expressions were restored in placenta and trophoblast cells after Mito-T.

**Interpretation:** This is the first study of PE treatment with Mito-T. Mito-T reverses pathological phenotypes of PE rats by improving placental mitochondrial and vascular function. The results provide proofs of concept of Mito-T as a potential therapeutic strategy for reducing maternal and fetal risks in PE patients.

**Funding:** This work is supported by Korean National Research Foundation of Korea (NRF) grant funded by the Korea government (MSIT) (NRF-2019R1A2C1005720, NRF-2023R1A2C1005720), BK21 FOUR education program, Korean Society of Hypertension (Grant number KSH-R-2020), National Natural Science Foundation of China (NSFC 31660284, NSFC31860288).

## 1. INTRODUCTION

Preeclampsia (PE) affects 2-8% of pregnant women worldwide, it is a leading cause of neonatal and maternal morbidity and mortality. ^1,2^ Clinical manifestations of PE include elevated blood pressure and kidney dysfunction (proteinuria) 20 weeks into the gestation. If leave untreated, maternal systemic responses develop to HELLP syndrome, pulmonary edema, seizure, and fatal arrhythmias which affects both mothers and babies.^3^ Currently, pre-term delivery and intravenous MgSO_4_ to relieve vasospasm are the only treatments for PE.^4^ Early effective strategies are urgently needed to prevent maternal and fetal death and abnormality.

The etiology of PE remains unclear. General consensus is that placental-originated ischemia and multiple organ dysfunction are the core to placental pathology of PE patients.^5,6,7^ Initially, insufficient trophoblast invasion and abnormal spiral arterial formation reduce uteroplacental perfusion, which exacerbates angiogenesis and placental ischemia. ^6^ Recent evidences suggest that the soluble fms-like tyrosine kinase 1(sFlt-1) levels are increased in the plasma and in placenta of PE patients, ^8,9^ The elevation of sFlt-1 and sFlt-1 to placental growth factor (PlGF) ratio in the sera of PE patients have been recognized as diagnostic biomarkers of PE.^9–12^ *In vitro* and *in vivo* studies show convincing evidences that hypoxia increases sFLT-1 secretion from trophoblast cells, sFLT-1 reduces vascular endothelial growth factor and plays primary roles in the development of maternal hypertension and abnormal spiral artery remodeling.^12^ In PE and PE-related patients, high level of sFLT-1 and/or sFLT-1/PlGF is strongly associated with adverse outcome (HELLP syndrome, pre-term delivery and low birth weight, etc.).^11^ Reduced blood supply disturbs metabolic status in the placenta ^13^ and mitochondrial dysfunction is critical to the angiogenesis and placental development. ^14,15^ Mechanistically, ischemia/hypoxia reduces mitochondrial content and mitochondrial oxidative capacity, leading to an increase in the production of reactive oxygen species (ROS) and oxidative stress.^15–17^

Currently, mitochondria transplantation (Mito-T) is emerging as a new therapy for ischemia-related diseases.^18^ Local and intravascular injection of autologous as well as heterologous mitochondria are proven to be beneficial to the patients of myocardial infarction, pre-ischemia, immediately after ischemia and during reperfusion.^19^ Application of Mito-T expends from diseases of the heart and brain (stroke) to kidney, lung, liver and skeletal muscle, spinal cord and sepsis.^19^ The source of mitochondria includes skeletal muscle, liver, heart and platelets. ^19^ The efficacy of mitochondria from various sources is demonstrated to be comparable, but more stable sources can be advantageous. Recently, mitochondria from human umbilical cord mesenchymal stem cells (hUC-MSC) have been shown to be integrative and functionally stable^20^, Mito-T from hUC-MSC improves the survival of LPS-injected sepsis in mice^20^. Phase I/IIa clinical trial study for refractory polymyositis/dermatomyositis indicates therapeutic potential, opens the possibility of hUC-MSC-derived mitochondria as a useful source for a wide range of human diseases. Until recently, Mito-T of hUC-MSC for placental dysfunction and PE has not been studied yet.

Here, the effects of Mito-T from hUC-MSC were tested in Ang II-induced PE rat model. Maternal and fetal phenotypes and vascular structure of maternal main organs were observed before and after Mito-T. In particular, placenta structure, mitochondrial function and dynamics were examined together with molecular mechanisms mediating functional recovery of placental mitochondria and vasculature of PE. Our results provide convincing evidence to indicate beneficial effects of Mito-T on PE maternal organs and placental mitochondrial function.

## 2 MATERIALS AND METHODS

### 2.1 Rat model of PE and Mito-T administration

Animal experimental protocols were performed in accordance with the Guide for the Care and Use of Laboratory Animals approved by the Institutional Animal Care and Use Committee of the Laboratory Animal Center, Seoul Medical University, South Korea [SNU-220724-1-1]. Twenty-four Sprague-Dawley (SD) pregnant rats (gestation day, GD 7, KOATECH, Korea) were randomly divided into four groups: sham, Ang II, sham+MT, Ang II+MT. Ang II was infused through osmotic minipump (1 µg/kg/min) at GD 8 in Ang II groups. hUC-MSC mitochondria (100 μg/μl) were injected into jugular vein (Mito-T groups) on GD 14 and rats were sacrificed at GD 20 (Figure 1a). Kidney and placenta tissues were collected at the day of sacrifice. Blood pressure was measured on GD 7, 9, 11, 13, 15, 17, 19, and 20 days using a Noninvasive Blood Pressure Measurement Device (CODA), averaged data from 3 cycles of measurements were collected for data analysis.

**Figure 1.**
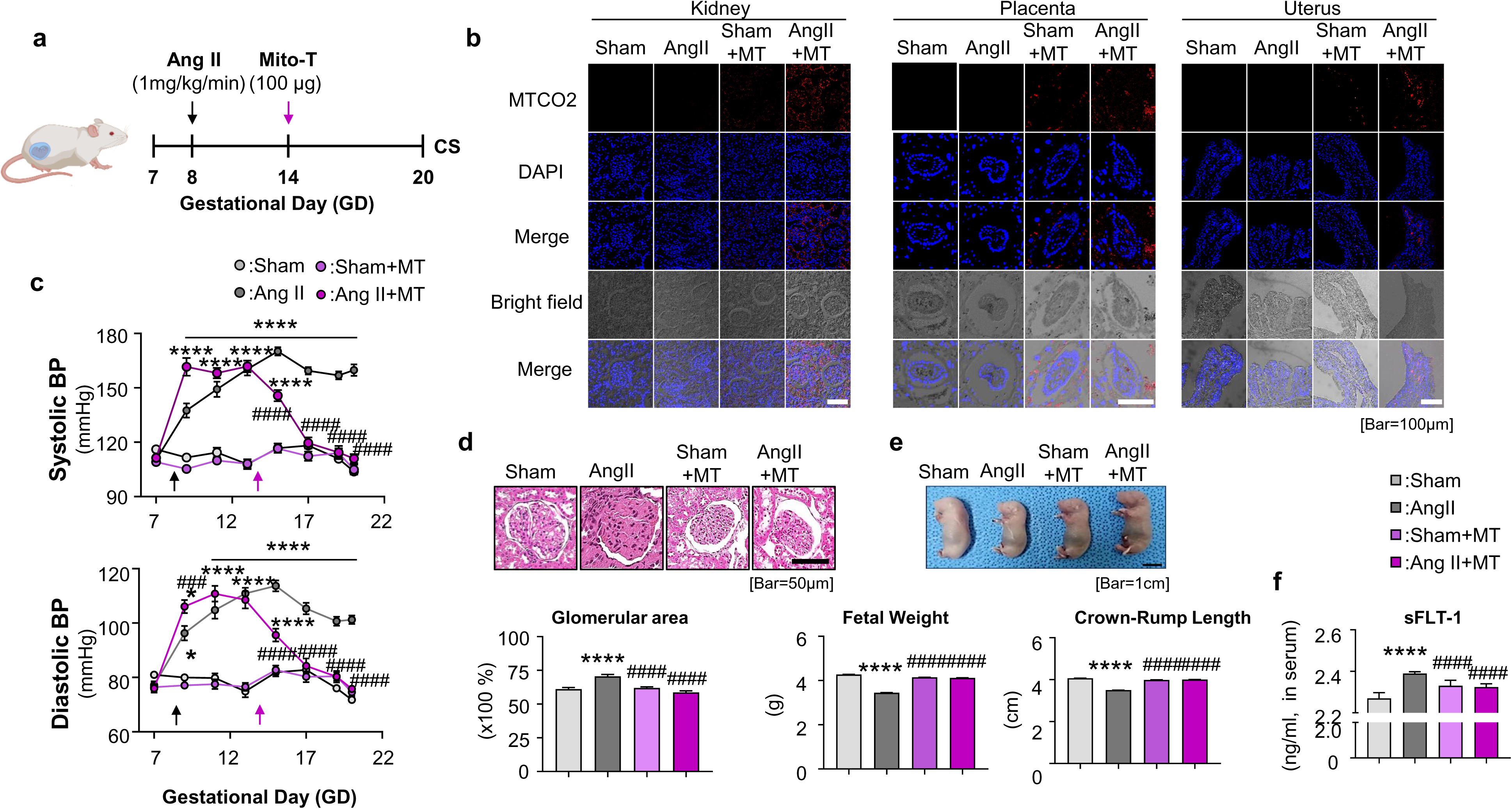
Mitochondria transplantation restores the clinical manifestations in Ang II-Induced Preeclampsia Rat Model. (a), Ang II was infused through osmotic minipump (1 µg/kg/min) at GD 8 in Ang II groups. hUC-MSC mitochondria (100 μg/μl) were injected into jugular vein (Mito-T groups) on GD 14 and rats were sacrificed at GD 20. (b), The distribution of hUC-MSC Mito in the kidney, uterus and placenta tissues of rats in Sham and Ang II groups. (c), Comparison of systolic and diastolic blood pressure in Sham, Ang II, Sham+MT, and Ang II+MT groups. (d), Representative H&E staining images of the kidney. Glomerular area was compared in 4 groups. (e), Images of fetal rats from Sham, Ang II, Sham+MT, Ang II+MT four groups. Fetal weight and fetal crown-rump length were compared among four groups. (f), Serum level of sFlt-1 in Sham, Ang II, Sham+MT, Ang II+MT four groups. Values are expressed as mean ± standard deviation range. *P<0.05, **P<0.005, ****P<0.0001

### 2.2 Isolation of mitochondria from hUC-MSC

hUC-MSC of passage 7 was used for mitochondria preparation. Briefly, hUC-MSC cells were harvested and depressurized in SHE buffer (0.25 M Sucrose, 20 mM HEPES, 2 mM EGTA, 0.1% bovine serum albumin, pH 7.4) using nitrogen cavitation (Parr Instrument, USA). After centrifugations (2,000 × g for 10 min at 4°C and the supernatant at 12,000 × g for 15 min, 4°C), the mitochondria pellet was washed twice in 500 μl SHE buffer followed by centrifugation at 20,000 × g for 10 min at 4°C. Resuspend the purified mitochondrial pellet in 100 μl suspending buffer and kept on ice until use. Protein concentrations were quantified using a bicinchoninic acid (BCA) assay.

### 2.3 H&E staining & immunohistochemistry

Fix the kidney and placenta tissues in 4% paraformaldehyde, wax sealed, slice cut, and dyed. Subsequently, the tissues were transferred to 70% ethanol solution for 24 hr at 20-22 °C. The samples were embedded in paraffin and cut into 4 µm sections, stained with hematoxylin/eosin (H&E) and toluidine blue. The slides were visualized using Aperio ImageScope software.

Paraffin-embedded tissue sections were rehydrated, demasked (using tris-EDTA (pH 8.0) and 10 mM sodium citrate buffer (pH 6.0) at 95°C for 10 minutes), blocked (with goat serum for 20 min) and was incubated with primary antibody (eNOS; 1:200, BD Biosiences) at 4°C followed by incubation with secondary antibody (Anti-IgG: 1:200, Cell Signaling) at room temperature for 30 min. The samples were processed using a DAB substrate kit (Sigma, Aldrich, USA), according to the manufacturer’s instructions. To track Mito-T, rehydrated tissue was stained using an antibody for human mitochondria cytochrome c oxidase subunit II (MTCO2; 1:1000, rabbit polyclonal antibody, Abcam). Wash with PBS and stain fluorescence-tagged rabbit secondary antibodies (Invitrogen, USA) for visualization. Stained samples were mounted in VECTASHIELD with DAPI mounting solution and imaged using the confocal microscope (Fluoview FV3000, Olympus, Japan).

### 2.4 ELISA

sFLT-1 levels in maternal serum samples were determined using enzyme-linked immunosorbent assay (ELISA) kits by MyBiosource Inc. (San Diego, CA, USA; CAT#MBS725733) according to the manufacturer’s instructions.

### 2.5 RNA preparation and quantitative real-time PCR

Total RNA was extracted from collected tissues using Qiazol Lysis Reagent (Cat.no.79306, QIAGEN). The purified RNA concentration was determined with NanoDrop (ThermoFisher). Primer information were shown below (Table 1). Briefly, cDNA was amplified with SYBR green (BIONEER 2XGreenStarqPCRMaster MIX) using a real-time PCR system (Applied Biosystems 7500) under the condition (95℃ for 3 min, 40 cycles at 95℃ for 10s, 55℃ for 3s, 95℃ for 10 s and 65℃ for 5s). Specificity of amplification products was assessed by melting curve analysis. Relative gene expression was calculated using the comparative threshold (Ct) method (2^-△△Ct^).

**Table 1.**
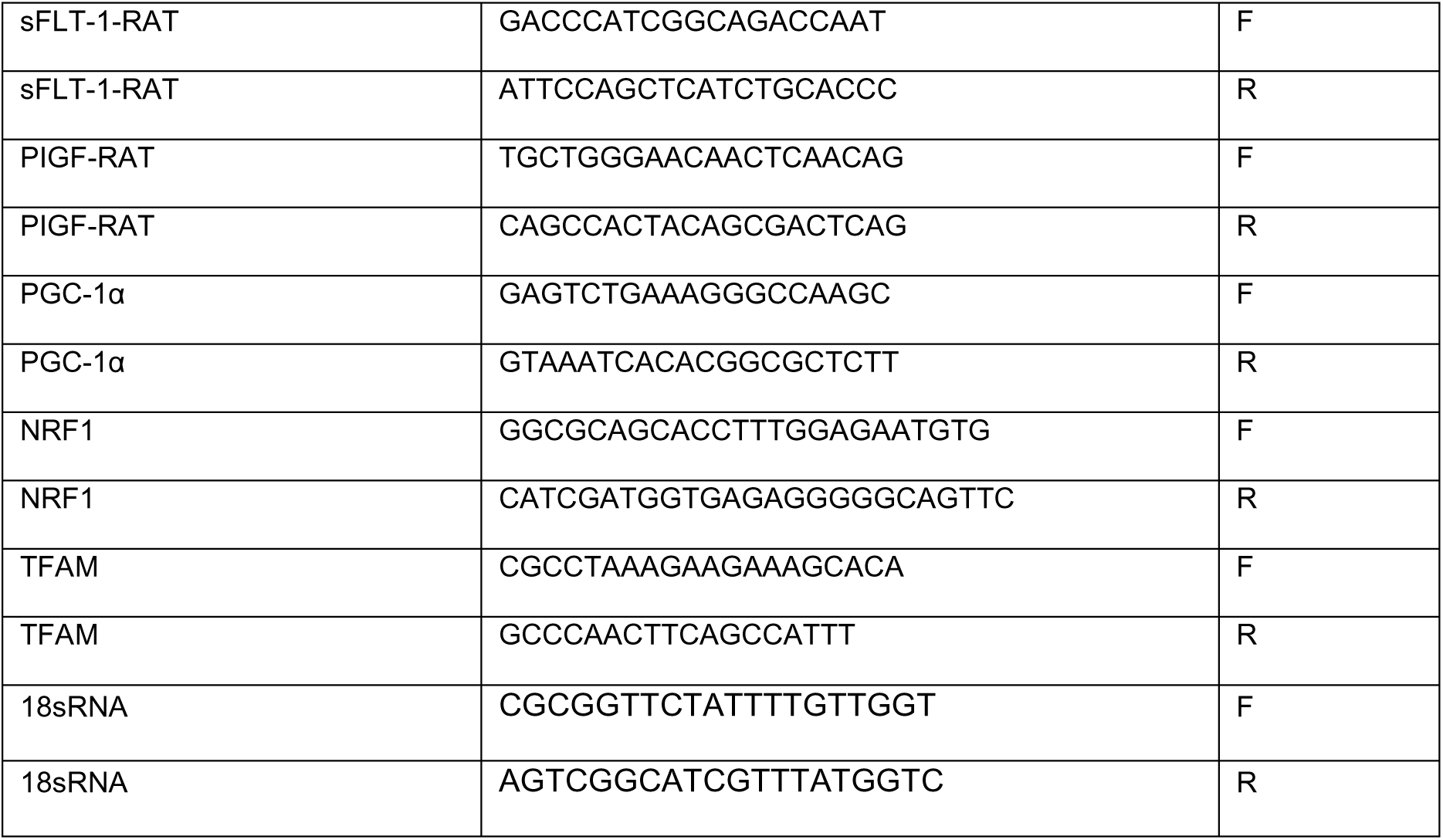
Primer sequences used in quantitative real-time PCR.

### 2.6 Western blotting analysis

Proteins were extracted from placenta tissues in lysis buffer containing 150mM NaCl, 50mM Tris-HCl, 1mM EDTA, 1% Triton X-100 with phosphatase inhibitor cocktail (Roche), pH 7.4. The lysates were boiled at 95℃ for 10 min and were fractionated by SDS/PAGE and transferred to PVDF membranes in 25mM Tris, 192mM glycine, 0.01% SDS, 20% methanol. Membranes were blocked in 1X TBS containing 1% Tween-20 and 5% BSA (blocking solution) for 1 hr at RT with gentle rocking. Membranes were incubated overnight at 4℃ with primary antibodies followed by secondary antibodies (Table 2). Blots were developed using ECL plus western blotting detection reagents (Amersham Bioscience).

**Table 2.**
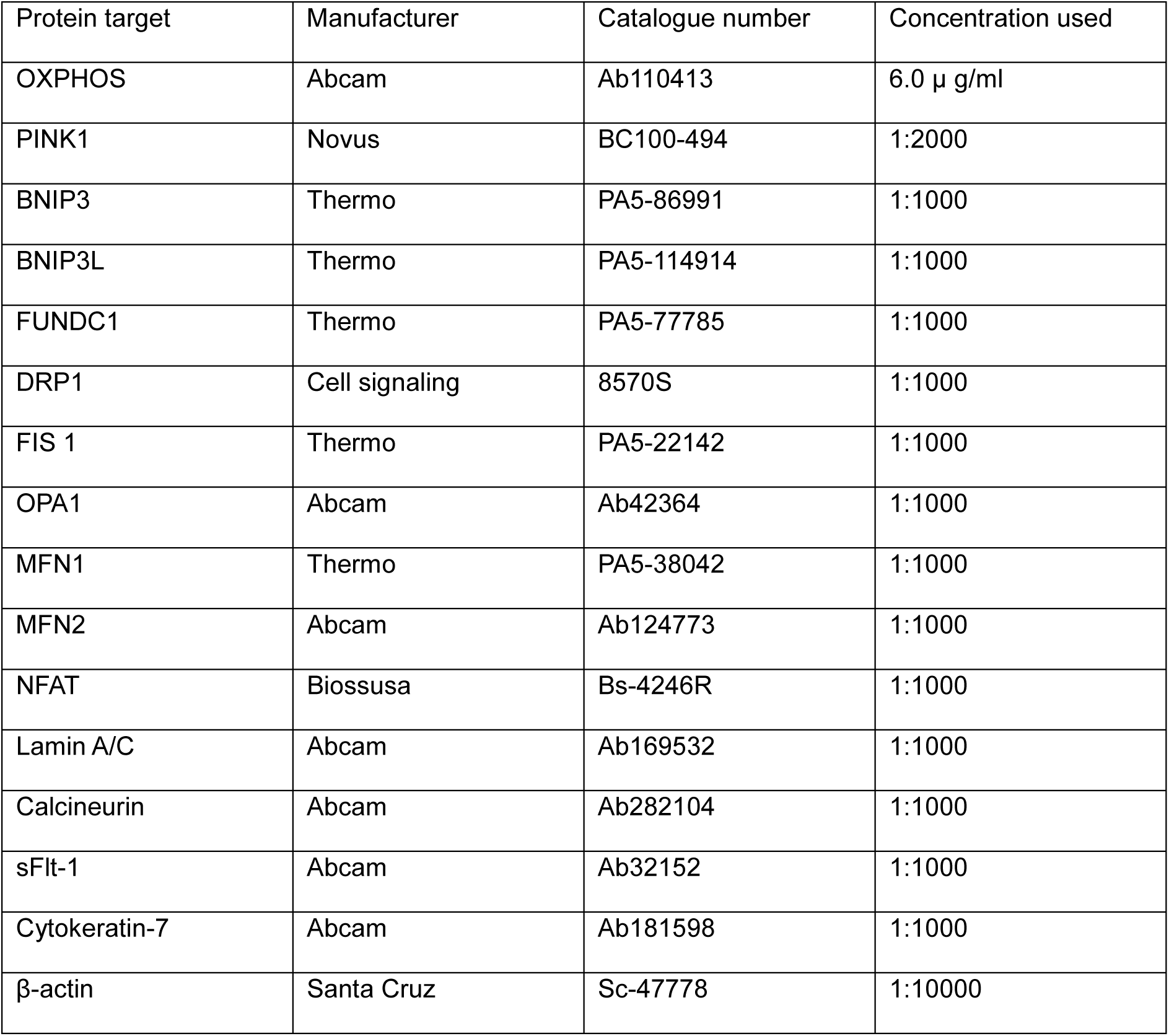
Primary antibodies used in western blotting.

| Protein target | Manufacturer | Catalogue number | Concentration used |
| --- | --- | --- | --- |
| OXPHOS | Abcam | Ab110413 | 6.0 µg/ml |
| PINK1 | Novus | BC100-494 | 1:2000 |
| BNIP3 | Thermo | PA5-86991 | 1:1000 |
| BNIP3L | Thermo | PA5-114914 | 1:1000 |
| FUNDC1 | Thermo | PA5-77785 | 1:1000 |
| DRP1 | Cell signaling | 8570S | 1:1000 |
| FIS 1 | Thermo | PA5-22142 | 1:1000 |
| OPA1 | Abcam | Ab42364 | 1:1000 |
| MFN1 | Thermo | PA5-38042 | 1:1000 |
| MFN2 | Abcam | Ab124773 | 1:1000 |
| NFAT | Biossusa | Bs-4246R | 1:1000 |
| Lamin A/C | Abcam | Ab169532 | 1:1000 |
| Calcineurin | Abcam | Ab282104 | 1:1000 |
| sFlt-1 | Abcam | Ab32152 | 1:1000 |
| Cytokeratin-7 | Abcam | Ab181598 | 1:1000 |
| β-actin | Santa Cruz | Sc-47778 | 1:10000 |

### 2.7 ATP synthesis activity assay

For measurement of ATP synthesis, reaction mixture containing 500μM ADP and 5μg mitochondria were added at room temperature and ATP was measured every 2.5 s for 20 min on a microplate reader at luminescent signal using SYNERGY HTX multi-mode reader (BioTek, USA).

### 2.8 I Citric acid synthesis activity assay

Citrate synthase activities were assessed in isolated mitochondria as described previously.^21^ Briefly, after adding 2 μg mitochondria and 10 mM acetyl CoA, the reaction was started by addition of 5,5’-Dithiobis (2-nitrobenzoic acid) (DTNB) (Sigma) and the change of absorbance was recorded at 450 nm every 1 min for 20 min using SYNERGY HTX multi-mode reader (BioTek, USA).

### 2.9 Mitochondrial membrane potential (MMP) measurement

Incubate mitochondria (10µg) with Tetramethylrhodamine Methyl Ester (TMRM, 100nM) in mitochondrial assay solution (MAS buffer, in mM; D-Mannitol 220, Sucrose 70, KH_2_PO_4_ 10, MgCl_2_ 5, HEPES 2, EGTA 1, BSA 0.2% W/V, pH 7.2 with KOH) for 20 min at 28℃. MMP was measured with TMRM-incubated mitochondria in the presence of substrates (in mM; pyruvate 5, malate 5, glutamate 5, ADP 4, and succinate 5) in 96 well plate (excitation at 540nm and emission at 580nm). All procedures in plate setup, scanning and analysis were conducted according to the manufacturer’s protocols.

### 2.10 Reactive oxygen species measurement

Mitochondrial reactive oxygen species were determined by Amplex™ Red Hydrogen Peroxide/Peroxidase Assay Kit (Invitrogen, A22188). Prepare 100μl working solution containing 100μM Amplex® Red reagent and 0.2 U/mL HRP (horseradish peroxidase). Add mitochondria 10μl (1 μg/μl) with 100µl Amplex® Red reagent /HRP working solution and supplement MAS buffer (in mM; D-Mannitol 220, Sucrose 70, KH_2_PO_4_ 10, MgCl_2_ 5, HEPES 2, EGTA 1, BSA 0.2% W/V, pH 7.2 with KOH)

to 155µl, incubate in the dark (37℃, 10 minutes). Add 45μl substrates (in mM; pyruvate 5, malate 5, glutamate 5, ADP 4, and succinate 5) to mitochondria (155μl). ROS was measured in 96 well plate (excited at 530nm and emission at 590nm).

### 2.11 Isolation of trophoblast cell from placenta

Placenta tissue was cut into small pieces (2 x 2 mm) and was washed thoroughly with PBS (0-4°C) to clean the blood. Mince the tissue in DMEM media (5ml/g) and digest with 1 mg/ml Dispase II, 0.5 mg/ml collagenase I and 0.1 mg/ml DNase I at 37 °C for 20 min each cycle (repeat 3 times). Equal volume of DMEM, containing10 % FBS, 100 U/ml penicillin and 100 μg/ml streptomycin, was added to the tissue mixture to stop enzyme degradation. The mixture was filtered through 100 μm and 40 µm strainers, respectively, to remove tissue debris. Trophoblast cell suspension was collected and centrifuged at 350 g for 10 min at 4 °C. Trophoblast cell pellets were resuspended in 5 ml DMEM and were layered on top of a preformed Percoll gradient (65 %, 55 %, 50 %, 45 %, 35 %, 30 % and 25 %) and centrifuged at 30000 g at 4 °C for 30 min. The layer between the 45 % and 35 % (density 1.050-1.060 g/ml) was collected and trophoblast cells were resuspended in DMEM and centrifuged at 350 g at 4 °C for 10 min. The pellets were suspended in PBS solution for further experiments.

### 2.12 Nuclear protein extraction

Nuclear protein extraction was performed by using nuclear extraction kit (Lot: WB 317789, Thermo scientific, USA) according to the manufacturer instruction. Briefly, 20 mg of placental tissue in 200 μl ice-cold buffer A (with DTT and protease inhibitor) was homogenized and centrifuged at 16000 x g for 10 min at 4°C. Buffer B (11 μl) was added to the pellets and the mixture was centrifuged at 16000 × g for 5 min at 4°C. The supernatant is taken as cytosol fraction of placental cells. The pellets were put on ice for 40 min after adding 100μl buffer C (with DTT and protease inhibitor) (vortex for 15 sec every 10 min). The mixture was then centrifuged at 16000 × g for 10 min at 4℃ and the supernatant was taken as nuclear fraction of placental cells.

### 2.13 Culture of trophoblast BeWo cell line

Trophoblast cell line BeWo was obtained from the Korean Cell Line Bank (KCLB). (KCLB No. 10098) BeWo cells were maintained in DMEM media (Thermo Fisher) supplemented with fetal bovine serum (FBS, 10%). The cell lines were separated into four groups: Sham, Sham+MT (100µg/ml) and Ang II (1µM), Ang II+MT. Following 24-hour incubation, cells were collected, sFLT-1, Calcinurine, protein expressions and eNOS, nNOS AT2R mRNA levels were investigated.

### 2.14 Statistical analysis

Statistical analyses were performed on GraphPad Prism 8 (GraphPad Software). All data are shown as means ± SEM or median (minimum to maximum). One-way ANOVA or independent t-test were used for statistical analysis. P<0.05 was considered a statistically significant difference. All experiments were repeated thrice.

## 3 Results

### 3.1 Mito-T restores clinical manifestation in Ang II-induced PE rats

First, we observed Mito-T distribution in sham and Ang II rat organs on GD20 through immunofluorescent staining of MTCO2 proteins of hUC-MSC. As shown in Figure 1b, Mito-T was distributed in the glomerular and collecting tubules of kidney, epithelial interstitial spaces of uterus and labyrinth/fetal vessels of placenta of both Sham and Ang II rats, with noticeably higher intensity in Ang II groups.

Figure 1c showed that Ang II infusion from GD8 caused significant increases in systolic and diastolic blood pressures (BP, mmHg) from GD9 to GD20 (peak BP in Sham: systolic BP 112.28 ± 12.72, diastolic BP 81.86 ± 8.95; Ang II: systolic BP 156.75 ± 11.58, diastolic BP 113.69 ± 12.06, Sham vs. Ang II P<0.0001, P<0.0001). Mito-T administration at GD14 gradually reduced BP of Ang II rats to GD20 without changing BP in Sham (Sham+MT: systolic BP 102.40 ± 12.51, diastolic BP 74.57 ± 7.67; Ang II+MT: systolic BP 107.47 ± 11.15, diastolic BP 75.80 ± 6.39, Sham vs. Ang II P<0.0001, Ang II vs. Ang II+MT P<0.0001). Across GD15-GD20, BP was significantly higher in Ang II compared to those in Ang II+MT (systolic BP, P<0.0001, diastolic BP, P<0.0001).

Next, we examined histological changes of maternal kidney cortex with Ang II and the effects of Mito-T were observed. As shown in Figure 1d, H&E results showed that glomerular capillaries were reduced and glomerular diameter was increased in Ang II group, so the proportion of glomerular diameter/capsule volume was significantly greater in this group (glomerular diameter/capsule volume: 61.12% in sham, 70.58% in Ang II, P<0.0001, Figure 1d). Mito-T restored glomerular capillaries and diameter in Ang II group and the proportion of glomerular diameter/capsule volume (61.92% in sham, 58.66% in Ang II, P<0.0001, Figure 1d), indicating the recovery of glomerular structure post-Mito-T.

The offspring of 4 groups were examined to investigate the outcome following Mito-T. As shown in Figure 1e, fetal size appeared significantly smaller in Ang II, and Mito-T resumed normal size. Quantitatively, fetal weight was significantly reduced from Ang II rats, which was reversed by Mito-T (fetal weight in g: Sham: 4.74±0.95, n=144, Sham+MT: 4.4±0.72, n=157; Ang II: 3.58±0.66, n=149; Ang II+MT: 4.16±0.64, n=124; P<0.0001 between Sham vs. Ang II and P<0.0001 between Ang II vs. Ang+MT). Similarly, fetal crown-rump length was shorter in Ang II which was prevented by Mito-T treatment (fetal crown-rump length in cm: Sham: 4.07±0.05, n=138; Ang II: 3.50±0.03, n=140; Sham+MT: 3.99±0.05, n=118; Ang II+MT: 4.00±0.05, n=114, P<0.0001 between Sham *vs*. Ang II and P<0.0001 between Ang II *vs.* Ang II+MT).

sFLT-1 responses to Mito-T are essential to determine the efficacity of Mito-T on PE. As shown in Figure 1F, sFlt-1 was elevated in the serum of Ang II rats and Mito-T attenuated such an increment (sFLT-1 in Sham in ng/ml: 2.27±0.08, Ang II: 2.39±0.24, Sham+MT: 2.31±0.06, Ang II+MT: 2.32±0.04; Sham vs. Ang II, P<0.0001; Ang II vs. Ang II+MT: P=0.04).

These results presented direct evidence that Ang II induced PE phenotype in rats with clinical manifestations (maternal hypertension, kidney abnormality and sFLT-1 elevation, fetal growth restriction). Mito-T reversed all effects exhibiting preventive potentials for PE.

### 3.2 Mito-T administration on PE rat placenta

Mito-T distribution in the placenta and the positive outcome of PE offspring suggest their preventive roles. We went on and examined both structural and functional responses of placenta to Mito-T. As shown in Figure 2a , placenta size was smaller and placental weight was lower in Ang II which was reversed by Mito-T (Placental weight in g in Sham: 0.53±0.007g, n=105; Ang II: 0.47±0.008g, n=111; Sham+MT: 0.53±0.007g, n=98; Ang II+MT: 0.52±0.010g, n=78; Sham vs. Ang II, P<0.0001; Ang II vs. Ang II+MT: P< 0.0001). Morphologically, H&E and immunohistochemistry results showed that decidual basal layers were wider and sinusoid space in labyrinth zone was reduced in Ang II rat placenta. In addition, cross-sectional view showed that the villus in labyrinth zone was significantly widened, both structural changes were restored with Mito-T (Figure 2b). Furthermore, fetal vessel diameters in chorion were reduced (Sham in µm: 28.39±14.21, Ang II: 11.72±7.02, Sham+MT: 36.16±13.61, Ang II+MT: 32.42±12.62; Sham vs. Ang II, P<0.0001; Ang II vs. Ang II+MT: P< 0.0001), with noticeable reduced numbers of trophoblasts around the vessels (Figure 2c).

**Figure 2.**
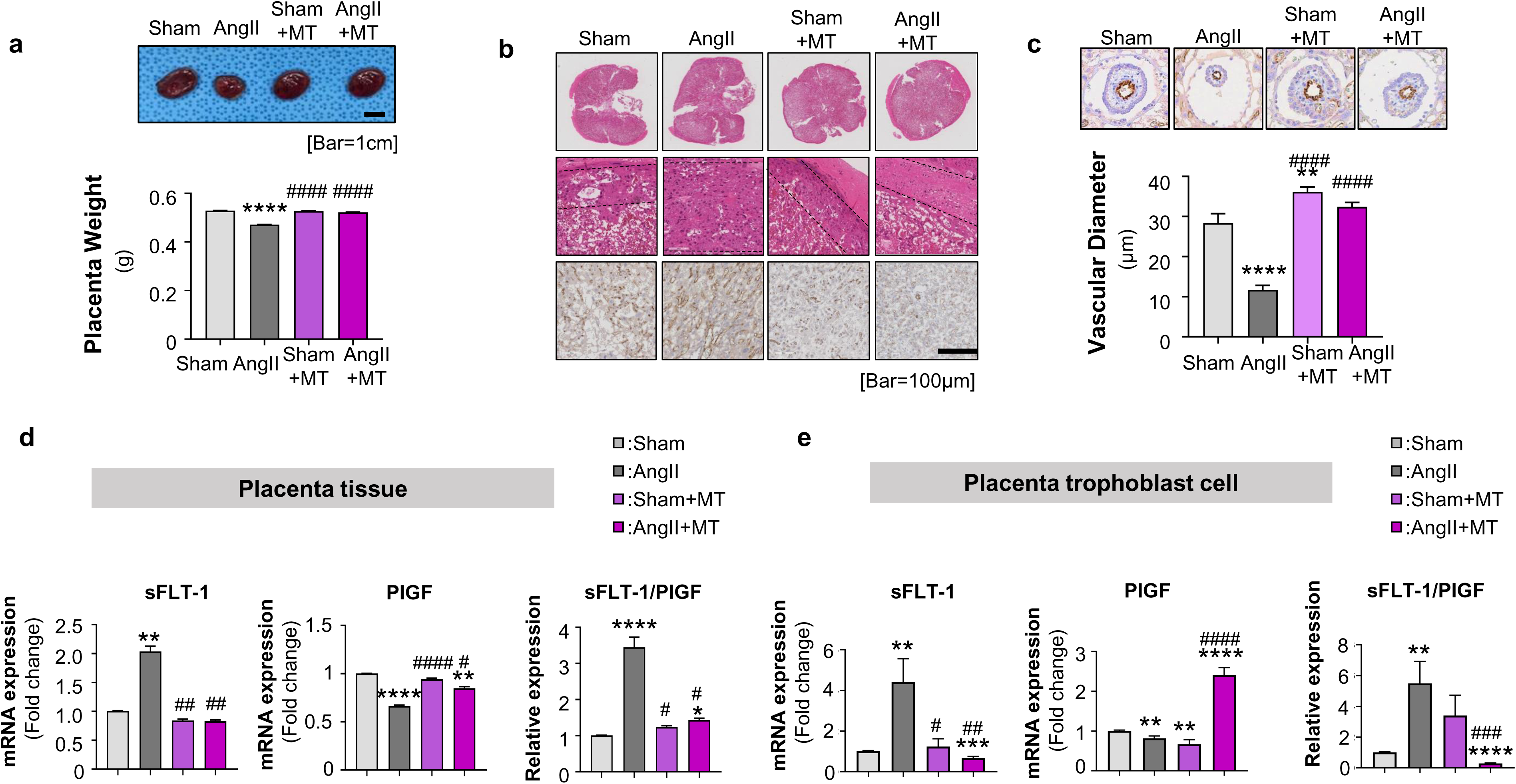
Mito-T reverses placental structure and sFLT-1 in Ang II-induced PE rats. (a), Representative images of placentas and the comparison of placental weight in four groups. (b). H&E and eNOS immunohistochemistry images showed changes in basal layer of the decidua, sinus space and the villi in the labyrinth zone, and fetal vessels. (c). Changes of placental chorionic fetal blood vessels in PE and Mito-T induced recovery. (d). sFlt-1, PlGF mRNA levels and sFlt-1/PlGF ratio in placental tissues of four groups. (e) sFlt-1, PlGF mRNA levels and sFlt-1/PlGF ratio in placental trophoblast cells of four groups. Values are expressed as mean ± SE. *P<0.05, **P<0.005, ****P<0.0001

Figure 2d showed that the mRNA expressions of sFLT-1 were significantly increased in Ang II which was reduced by Mito-T (tissue sFLT-1: Sham: 1.01±0.02, Ang II: 2.04±0.39, Sham+MT: 0.84±0.13, Ang II+MT: 0.83±0.12; Sham vs. Ang II, P<0.0001; Ang II vs. Ang II+MT: P<0.0001). Another important pro-angiogenesis factor, PlGF was examined and the results showed reduced PlGF mRNA in Ang II group which was reversed by Mito-T (tissue PlGF: Sham: 1.00±0.010, Ang II: 0.66±0.05, Sham+MT: 0.94±0.05, Ang II+MT: 0.85±0.07; Sham vs. Ang II, P<0.0001; Ang II vs. Ang II+MT: P< 0.0001). Consequently, sFlt-1/PlGF ratio, an important placenta biomarker, was increased in Ang II placenta but Mito-T attenuated such an increment (sFLT-1/PlGF: 1.01±0.02, Ang II: 3.45±1.02, Sham+MT: 1.24±0.16, Ang II+MT: 1.43±0.21; Sham vs. Ang II, P<0.0001; Ang II vs. Ang II+MT: P< 0.0001, Figure 2d). Similar results were shown in isolated trophoblast cells. To confirm, sFLT-1 and PlGF were observed in isolated trophoblast cells. Figure 2E showed that sFLT-1 mRNA expression was greater in Ang II compared to sham (sFLT-1: Sham: 1.01±0.14, Ang II: 4.41±4.87, p=0.0055) and MitoT reduced sFLT-1 level (Sham+MT: 1.24±1.63, Ang II+MT: 0.67±0.37; Ang II vs. Ang II+MT: P=0.0026). In addition, PlGF mRNA was reduced in Ang II (Sham: 1.00±0.09, Ang II: 0.82±0.24, p= 0.0040) and Mito T increased it (Sham+MT: 0.67±0.48, Ang II+MT: 0.21±0.80; Ang II vs. Ang II+MT: P< 0.0001). As a result, sFlt-1:PlGF ratio was significantly high in Ang II (Sham: 1.01±0.15, Ang II: 5.49±6.04, Sham vs. Ang II, P=0.0034) and Mito-T reduced it (Sham+MT: 3.40±5.62, Ang II+MT: 0.29±0.14; Ang II vs. Ang II+MT: P=0.0009, Figure 2e).

These results demonstrate that Mito-T reverses placental structures and functions, through reducing sFLT-1 and increasing PlGF mRNA levels in PE.

### 3.3 Mito-T on placental mitochondria functions

Preventive effects of Mito-T suggest an improvement of placental mitochondria functions. First, we examined the protein expressions of key components of electron transport chain in the placenta of 4 groups. As shown in Figure 3a&b, complex I, II subunits were significantly reduced in Ang II group (Sham vs. Ang II CI P=0.0267, CII P=0.0035) but Mito-T increased all complex I-V components (Ang II *vs.* Ang II+MT CI P=0.0250, CII P<0.0001, CIII P=0.0013, CIV P=0.0005, CV P=0.025 Figure 3 a&b)

**Figure 3.**
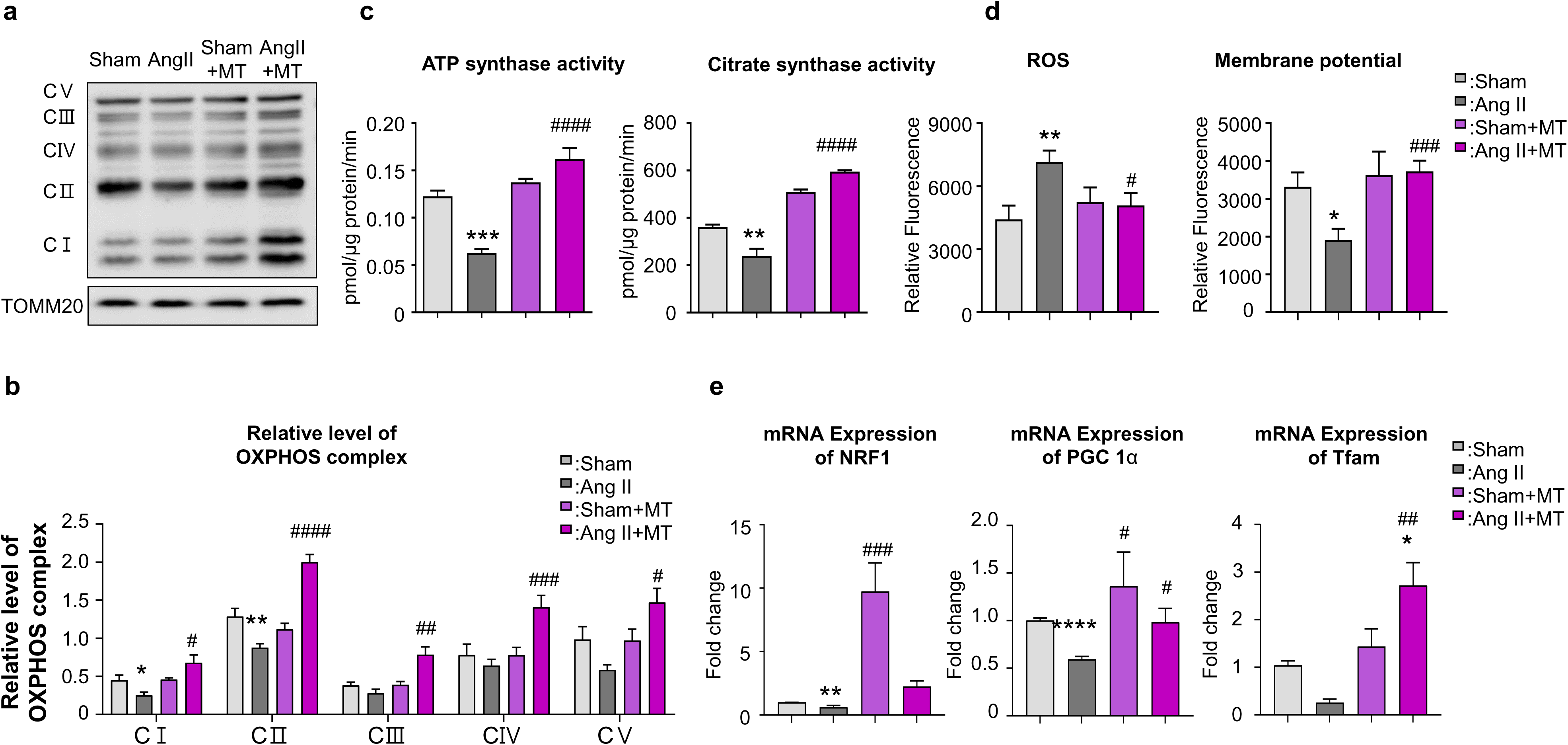
Effects of Mito T on mitochondrial contents and activity in PE rat placenta. (a & b). Comparison of mitochondrial complexes in Sham, Ang II, Sham+MT, Ang II+MT. (c&d) Functional aspects of mitochondrial activities in the placenta of 4 groups (ATP synthase activity and citrate synthase activity; Mitochondrial ROS and MMP). (e), mRNA expressions of mitochondrial biogenesis markers in four groups (NRF1, PGC 1, TFAM). Averaged results were expressed as mean ± SE. *P<0.05, **P<0.005, ****P<0.0001

Both mitochondrial ATPase and citrate synthase activities were reduced in Ang II, and Mito-T significantly increased the activities (ATP in pmol/ μg protein/min: Sham *vs.* Ang II: P=0.0002; Ang II *vs.* Ang II+MT P<0.0001; Citrate in pmol/ μg protein/min: Sham *vs.* Ang II: P=0.0085; Ang II *vs.* Ang II+MT P<0.0001 Figure 3c). Importantly, placental ROS was increased and mitochondrial membrane potential was reduced in Ang II (ROS: Sham *vs.* Ang II: P=0.0059; MMP: Sham *vs.* Ang II: P=0.0059, Figure 3d), but Mito-T reversed all changes (ROS: Ang II *vs.* Ang II+MT P=0.0217; MMP: Ang II *vs.* Ang II+MT P=0.001, Figure 3d).

In addition, Figure 3e showed that mRNA of mitochondrial biogenesis markers, peroxisome proliferator-activated receptor gamma coactivator 1α (PGC-1α), nuclear respiratory factor (NRF1) and mitochondrial transcription factor (TFAM) expression were reduced in Ang II, but Mito-T increased the levels of PGC-1a, NRF1 and TFAM (Sham vs. Ang II: PGC-1 α P<0.0001, NRF1 P=0.0131, TFAM P<0.0001; Ang II vs. Sham+MT : PGC-1 α P=0.0484, NRF1 P=0.0005, TFAM P=0.0054 ; Ang II vs. Ang II+MT: PGC-1 α P=0.0189, NRF1 P=0.0017, TFAM P<0.0001).

### 3.4 Mito-T on placental mitochondrial dynamics and mitophagy biomarkers

Mito T may improve mitochondrial function through affecting placental mitochondrial dynamics. Representative mitochondrial fission protein, dynamin related protein 1(DRP1) was shown to be increased and Fission 1(Fis1) was reduced in Ang II rats but Mito-T reversed these changes (DRP1: Sham vs. Ang II: P=0.0249; Ang II vs. Ang II+MT: DRP1 P<0.0001; Fis 1: Sham vs. Ang II: P<0.0001, Ang II vs. Ang II+MT: P=0.0035, Figure 4a)

**Figure 4.**
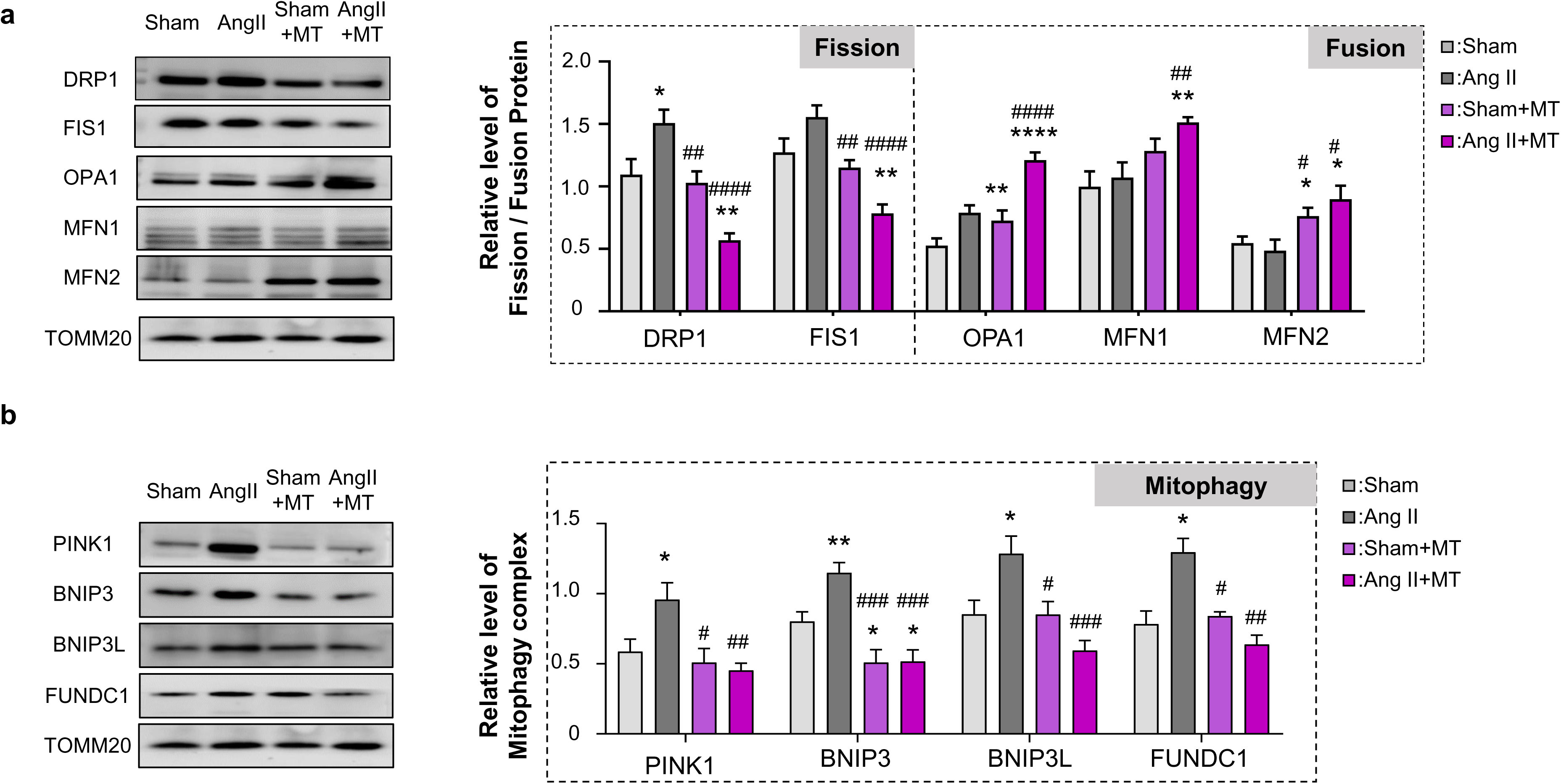
Mitochondria dynamics are altered – fission is reduced but fusion is improved following Mito-T. Mitophagy was reduced in PE rat placental mitochondria. (a), Comparison of mitochondrial fission & fusion proteins (fission proteins: DRP1, FIS1; fusion proteins: OPA1, MFM1, MFN2) (b),Comparison of mitophagy proteins (PINK1, BNIP3, BNIP3L, FUNDC 1) . Values are expressed as mean ± standard deviation range. *P<0.05, **P<0.005, ****P<0.0001

Mitochondrial fusion status was observed by detecting representative fusion protein expressions. As shown in Figure 4a, Optic atrophy 1 (OPA1) was increased in Ang II rat placenta (OPA1: Sham vs. Ang II, P=0.0071) but Mitofusin (MFN1 and MFN2) were not different between Sham and Ang II. However, OPA1, MFN1 and MFN2 protein expressions were increased and significant increment was observed in Ang II+MT group (Ang II vs. Ang II+MT: OPA1 P=0.0002, MFN1 P=0.0063, MFN2 P=0.0119).

Figure 4b showed that mitophagy biomarkers, PTEN induced putative kinase 1(PINK1), Bcl2/adenovirus E1B 19 kDa protein interacting protein 3 (BNIP3), BNIP3 like (BNIP3L), Fun14 domain containing 1 (FUNDC 1) were significantly increased in Ang II rat placenta of GD20 compared to those in Sham (Sham vs. Ang II, PINK 1 P=0.0325, BNIP3 P=0.0058, BNIP3L P=0.0196). Mito T significantly reduced these proteins (Ang II vs. Ang II+MT: PINK 1 P=0.0032, BNIP3 P=0.0002, BNIP3L P=0.0004, FUNDC1 P=0.0052, Figure 4b).

Taken together, these results suggest that Mito-T administration reduces placental mitochondria fission and improves fusion, mitophagy biomarkers are normalized with Mito-T at GD20, further suggest improved mitochondrial functions.

### 3.5 sFLT-1 and the regulating signaling pathways in placenta tissue, human Bewo cell line, trophoblast cells with and without Mito-T

Calcineurin and Nuclear factor of activated T-cell (NFAT) signaling is established to induce sFlt-1 elevation in PE placenta.^21,22^ As such, we examined calcineurin/NFAT pathways in the placenta of 4 groups to understand the mechanism of sFLT-1 reduction by Mito-T.

Figure 5a showed that sFLT 1 protein expression was significantly increased in Ang II rat placental tissue (Sham vs. Ang II, P=0.0229), which was reduced by Mito T (Ang II vs. Ang II+MT, P=0.0127). The protein levels of calcineurin was significantly increased in Ang II rat placenta which was reduced with Mito-T (Sham vs. Ang II: Calcineurin P=0.0054, Ang II vs. Ang II + mito: Calcineurin P=0.0003, Figure 5a). Further analysis showed that cytosol and nuclear NFAT protein level were significantly increased in Ang II rat placenta (cytosol P=0.0033, nuclear P=0.0493, Figure 5b), Mito-T reduced NFAT protein expression in the nucleus but maintained high expression level of NFAT protein in the cytosol (Ang II vs. Ang II + mito: NFAT cytosol P=0.0001, Figure 5b).

**Figure 5.**
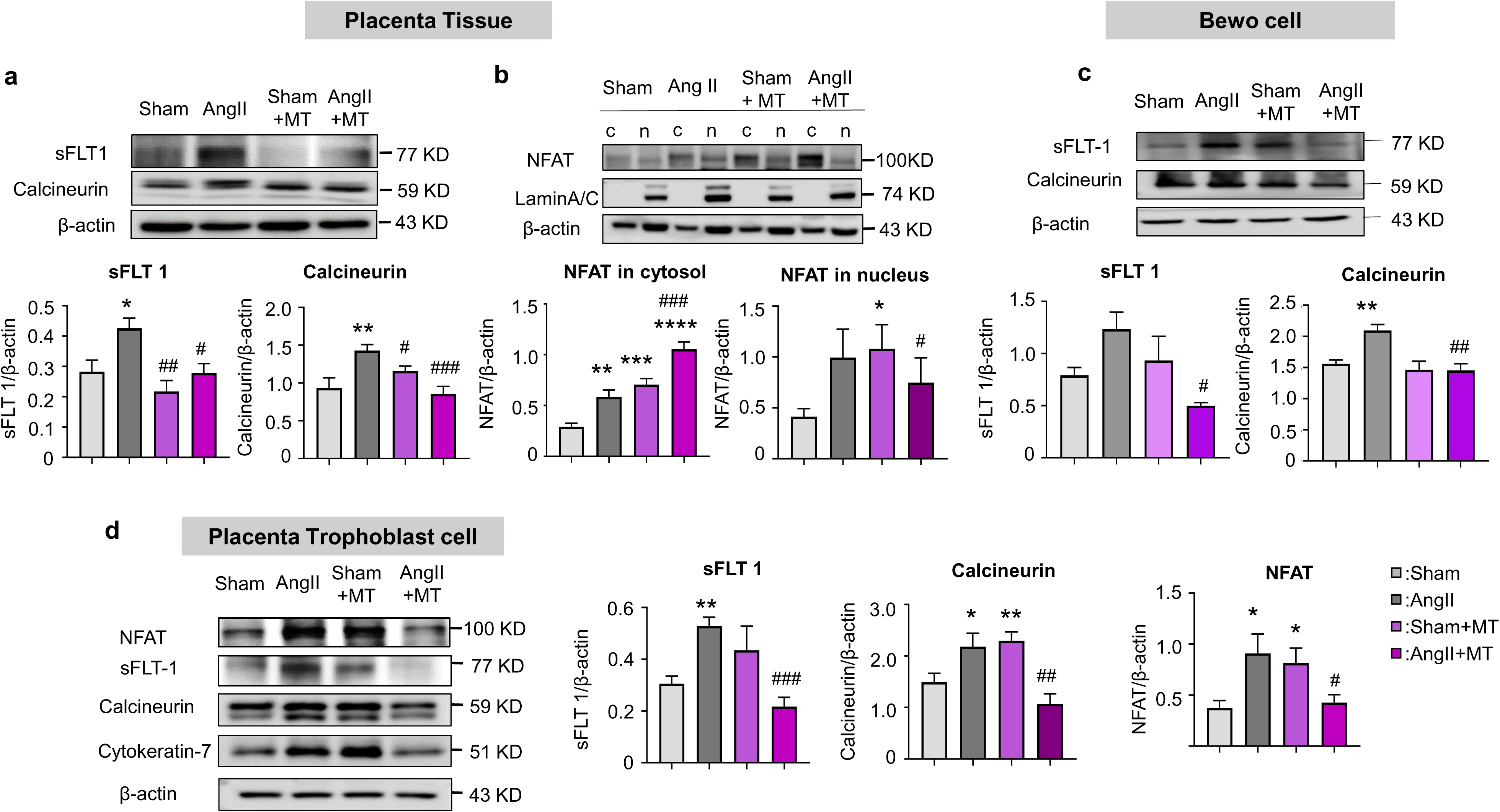
Calcineurin and NFAT signaling pathway with and without Mito-T in PE rat placenta. (a). sFlt-1 protein expression in placental tissue in four groups. (b). Calcineurin and NFAT protein expressions in the cytosol and nucleus of placental tissue. (c). Comparison of sFlt-1 protein levels and calcineurin in BeWo cell lines. (d). Protein expressions of calcineurin, NFAT and sFlt-1 in trophoblast cells isolated from four groups of placentas. Values are expressed as mean ± standard deviation range. *P<0.05, **P<0.005, ****P<0.0001.

Figure 5c showed that sFlt-1 protein expression was upregulated with Ang II treatment, which was significantly reduced following Mito-T treatment (Ang II vs. Ang II + mito, P=0.018, Figure 5c). In the same cell lines, protein expression of calcineurin was significantly increased with Ang II, which was reduced by Mito-T treatment (Sham vs. Ang II, P=0.0071, Ang II vs. Ang II + mito, P= 0.0012, Figure 5c).

The protein expressions of calcineurin, NFAT and sFLT-1 were also examined in isolated trophoblast cells. As shown in Figure 5d, calcineurin and NFAT protein expressions were increased in Ang II and Mito-T reduced it (Calcineurin: P=0.0496, NFAT 1 P=0.0299 between Sham and Ang II, Calcineurin P=0.0063 NFAT P=0.0007 between Ang II vs. Ang II + MT). In the same trophoblast cells of placenta, sFlt-1 was increased in Ang II (P=0.0025) and Mito-T reduced the increment (P=0.0007) (Figure 5d).

These results clearly demonstrate that sFLT-1 level was increased in placental trophoblast of Ang II rats and greater expression of calcineurin and NFAT nuclear translocation could be associated with sFLT-1 elevation. Mito-T reduced sFLT-1 in trophoblast cells through moderating calcineurin and NFAT signaling.

### 3.6 NOS signaling mediating vascular restoration in the trophoblast and placental tissue of Ang II rats with and without Mito-T

Immunohistochemistry results from Figure 6a showed that eNOS protein was expressed in the endothelial layer of placental vasculature in sham, sham+MT, Ang II and Ang II+MT groups. eNOS mRNA expression, which was significantly reduced in Ang II rat placenta, was reversed by Mito-T (eNOS: Sham vs. Ang II, P=0.0010; Sham+MT vs. Ang II + MT, P=0.0368, Figure 6a&d). nNOS protein expression was observed in trophoblast cells of the terminal villi in the placenta (Figure 6b). nNOS mRNA expression was reduced in Ang II rat placenta, which was reversed by Mito-T (nNOS: Sham vs. Ang II, P=0.0002; Sham+MT vs. Ang II + MT, P=0.0012, Figure 6b&c). Similarly, AT2R expression was observed in trophoblast cells of the terminal villi (AT2R: Sham vs. Ang II, P<0.0001; Sham+MT vs. Ang II + MT, P<0.0001, Figure 6c&f).

**Figure 6.**
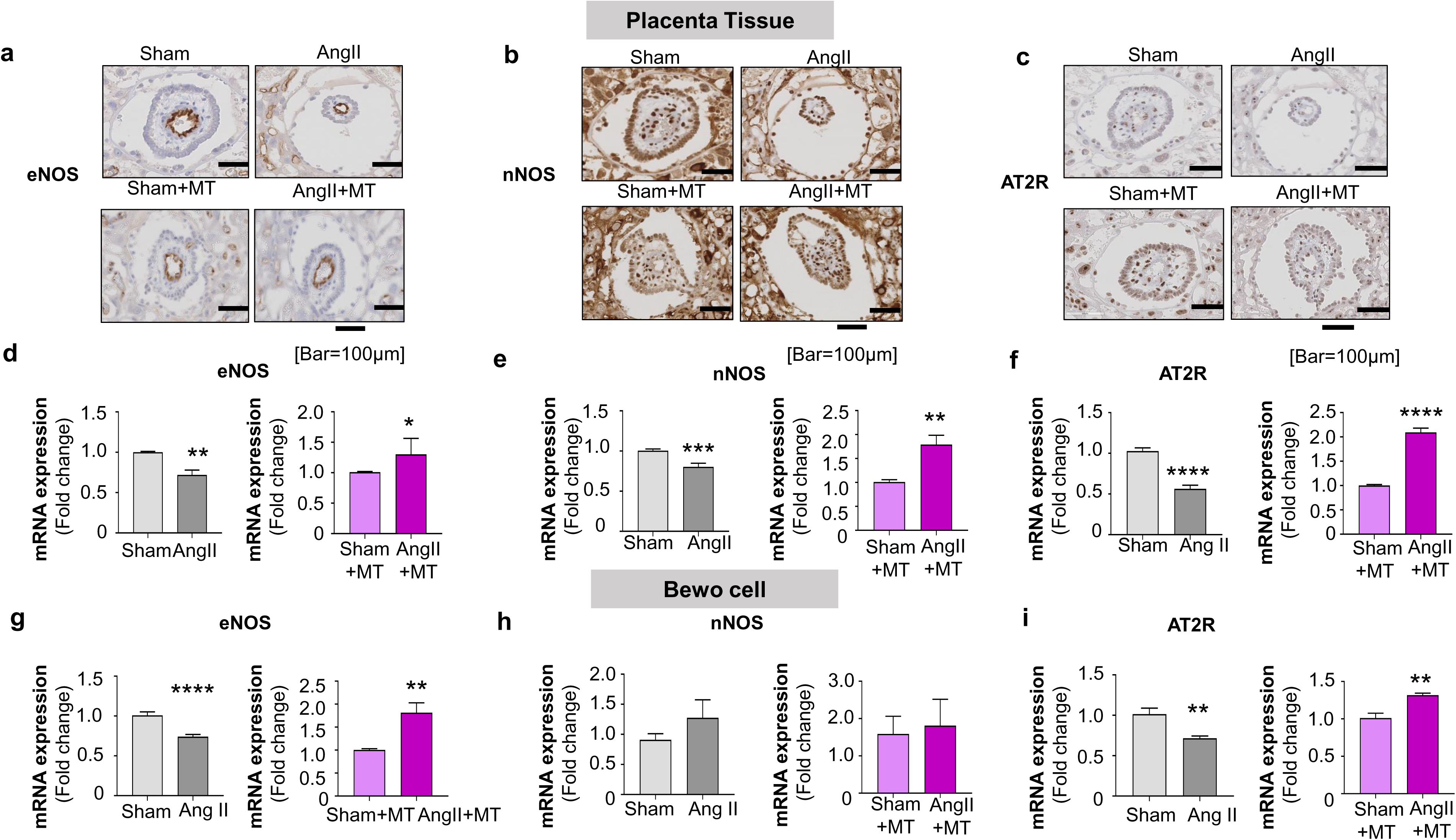
Localization and protein expression of AT2R, eNOS and nNOS in Sham, Ang II, Sham+MT, Ang II+MT rat placenta tissues ; mRNA expressions of AT2R, eNOS and nNOS in the BeWo cell line with and without Ang II and Mito-T. (a). IHC staining shows that eNOS protein is expressed on placental vascular endothelial cells. (d). Comparison of eNOS mRNA in placental tissues of the four groups. (b), IHC staining shows that nNOS protein is expressed on the trophoblast cells of the terminal villi of the placenta. (e), Compare the expression of nNOS mRNA in placental tissues of 4 groups. (c), IHC staining shows that AT2R protein is expressed on the trophoblast cells of the terminal villi of the placenta. (f), Comparison of AT2R mRNA expression in placental tissues of 4 groups. Values are expressed as mean ± standard deviation range. *P<0.05, **P<0.005, ****P<0.0001. (g ) , Comparison of eNOS mRNA expression in Sham, Ang II, Sham+Mito-T, Ang II+Mito T. (h), Comparison of nNOS mRNA expression in four groups. (i), Comparison of AT2R mRNA expression in four groups. Values are expressed as mean ± standard deviation range. *P<0.05, **P<0.005, ****P<0.0001.

Figure 6g-i showed the mRNA expressions of AT2R, eNOS, and nNOS in BeWo cell lines. Similarly to those observed in placental tissue, eNOS and AT2R mRNA expression, which were significantly decreased in the Ang II group, was increased after Mito-T administration (eNOS: Sham vs. Ang II, P<0.0001; Sham+MT vs. Ang II + MT, P=0.0018, Figure 6g; AT2R: Sham vs. Ang II, P=0.0033; Sham+MT vs. Ang II + MT, P=0.0015, Figure 6i). nNOS mRNA was reduced with Ang II (Figure 6h), and there was a trend towards an increase of nNOS mRNA in Ang II+MT (Figure 6h).

## 4 Discussion

The current study demonstrates that hUC MSC-derived mitochondria administration mid-term through PE can be protective against maternal and fetal abnormality. Experimental findings support the notion are: 1) Mito-T was able to control high blood pressure of PE rats and reverses the glomerulus structural dysregulation; 2) Mito-T increased fetal weight and crown rump length which were reduced in PE; 3) sFLT-1 mRNA and proteins were increased in maternal sera, placental tissue and trophoblast cells; Mito-T reduced sFLT-1 and sFLT-1/PlGF ratio, indicative of vascular normalization in the placenta. Indeed, histology examination demonstrated disease phenotypes in placental structure and fetal vessels of PE rats and Mito-T reversed all the changes to normal level; 4) Mito-T improved placental mitochondria functions (ATP synthase activity, citrate synthase activity, ROS reduction, MMP, biogenesis marker mRNA increment) and increased key elements of mitochondrial ETC complexes; 5) Mito-T administration resets the mitochondrial to fusion-status by reducing mitochondrial fission proteins and increasing fusion proteins, mitophagy was also normalized, indicative of improved mitochondrial stability; 6) In placental tissues, isolated trophoblast cells and cultured BeWo cell lines, sFLT-1, calcineurin and nuclear NFAT 1 proteins were increased in PE rats, Mito-T reduced the increment of these proteins. 7) Placental vascular normalization was accompanied by upregulation of eNOS, nNOS and AT2R with Mito-T. Here, we provide, *for the first time*, the prove of concept of the advantageous effects of hUC-MSC-derived mitochondria on PE.

Placental mal-perfusion impacts the maternal and fetal metabolism through mitochondrial dysregulation, including the reduction of the key components of ETC and increases in oxidative stress secondary to reduced oxidative capacity, consequently attenuating mitochondrial biogenesis ^22,23,24^. Our results clearly showed that Mito-T administration improved mitochondrial ETC functions (increase MMP, reduces ROS) and biogenesis (increase mRNA expressions of NRF, Tfam and PGC 1α). Protein expressions of complex I-V are increased suggesting mitochondrial DNA upregulation and post-translational functional regulations. Along the line, Mito-T reversed dysregulations of mitochondrial dynamics and mitophagy in PE, *i.e*. Mito-T administration reduced mitochondrial fission and mitophagy to normal levels and increased mitochondrial fusion in placental trophoblast cells. Exact nature of Mito T transformation in placenta of PE rats from GD14 to GD20 is unknown. Transplanted mitochondria either injected locally, or intravascular injection or incubation in culture media are found to be uptaken within minutes into various cell types of target organs through endocytosis.^25–29^ Specific labeling of transplanted mitochondria indicates that majority of mitochondria undergo integration with local mitochondrial network (∼80%), result in greater ATP synthesis and cellular ATP contents.^25^ In the current PE model, we observed that labeled hUC MSC mitochondrial proteins ^25,30–32^ were expressed in the kidney and placenta from GD15 till GD17 and GD20. The results that mitochondrial fusion is increased in PE with Mito-T speculate that transplanted hUC MSC mitochondria are either maintained through fusion processes in these tissue cells; alternatively, transplanted mitochondrial components could be produced through fusion with endogenous mitochondria and the replacement of mitochondrial DNA, inducing the translations and protein processes.^25,26,33^ This is essential for mitochondrial quality control those are damaged in PE with Mito-T during recovery stages.

The stability of placental trophoblast cells is prerequisite for sFLT-1 reduction and disease recovery in PE. Biochemical analysis indicated that calcinurine-NFAT -dependent signaling is upregulated, which was responsible for sFLT-1 increment in PE, whereas Mito-T reversed the pathways. Similar findings were confirmed in isolated trophoblast cells and human trophoblast cell line (BeWo cells). In Bewo cells, Ang II treatment for 24 hours significantly increased sFLT-1 production, which was associated with increased calcinurine in the cytosol. The expression of NFAT protein in BeWo cells with Ang II were variable and its nucleus translocation needs to be confirmed. Nevertheless, eNOS, nNOS and AT2R, which are well established to mediate protective effects in the vascular endothelial cells and trophoblast cells, were upregulated by Mito-T. Taken together, these results strongly support that hUC-MSC mitochondria can be a promising target for PE therapy drug development.

Accumulating evidences show consistent results with Mito-T to exert beneficial effects on various ischemic related diseases including PE.^34–42^ These effects could be through systemic as well as local regulations. E.g. immune responses of cross species mitochondria could exacerbating the inflammation and consequent cardiovascular events. However, Mito-T has not been associated with autoimmune or inflammatory reactions.^33,39,43–45^ In fact, recent experimental results with hUC MSC mitochondria on LPS induced sepsis mice showed attenuation of systemic inflammation.^46,47^ It is well established that hypertension or PE is associated with oxidative stress and inflammation,^48–56^ which affects systemic cardiovascular function, in particular kidney dysfunction.^57–61^ The facts that Mito-T recovered kidney structure, restored fetal weight and crown rump length at GD20, importantly, fetal vessel sizes were recovered with Mito-T, strongly suggest the safety of Mito T use without causing additional complications hampers fetus delivery.

Can Mito-T from hUC MSC be used in human PE? We consider it is possible because compelling evidence showed that injected hUC-MSC mitochondria were distributed at system organs up to 7 days post-transplantation and mitochondrial activity was upregulated and maintained up to the period of experiments. Beneficial effects of Mito-T on isolated heart, wound healing and inflammatory disease by improving mitochondrial activity. Safety approval was obtained with hUC-MSC derived mitochondria from toxicology and dose range finding tests. Mito-T can be a quick and effective regime for short- or long-term clinical research and disease therapy.

In Conclusion, Mito T is a novel regime for PE and hUC MSC origin of mitochondria provides stable and efficient potential for PE treatment. Our study using animal models of PE provided prove of concepts of Mito T application for this disease. In humans, systematic and thorough investigations of the dose and time of Mito T injection is needed to better define the blueprint for PE therapy.

## ACKNOWLEDGEMENT

This work is supported by Korean National Research Foundation of Korea (NRF) grant funded by the Korea government (MSIT) (NRF-2019R1A2C1005720, NRF-2023R1A2C1005720), BK21 FOUR education program, Korean Society of Hypertension (Grant number KSH-R-2020), National Natural Science Foundation of China (NSFC 31660284, NSFC31860288).

## DATA AVAILABILITY STATEMENT

All data are included in the manuscript or supporting materials.

## ETHICS STATEMENT

Animal experiments were performed in accordance with the Guide for the Care and Use of Laboratory Animals approved by the Institutional Animal Care and Use Committee of the Laboratory Animal Center, Seoul Medical University, South Korea [SNU-220724-1-1].

## Nonstandard Abbreviation and Acronyms

Ang II: Angiotensin II
Mito-T: Mitochondrial transplantation
sFlt-1: Soluble fms-like tyrosine kinase 1
PlGF: Placental growth factor
PE: Preeclampsia
GD: Gestational day
SD: Sprague-Dawley
BP: Blood pressure
ROS: Reactive oxygen species
hUC-MSC: Human umbilical cord mesenchymal stem cells
NFAT: Nuclear factor of activated T-cell
MMP: Mitochondrial membrane potential
H&E: Hematoxylin/eosin
PGC-1a: Peroxisome proliferator-activated receptor gamma coactivator 1-alpha
NRF1: Nuclear respiratory factor 1
Tfam: Mitochondrial transcription factor A
OPA1: Optic atrophy 1
DRP1: Dynamin-related protein 1
FIS1: Fission 1
MFN: Mitofusin
PINK1: PTEN-induced putative kinase 1
BNIP3: Bcl2/adenovirus E1B 19 kDa protein-interacting protein 3
BNIP3L: BNIP3-like
FUNDC1: Fun14 domain containing 1

## Graphical Abstract

**Figure 7.**
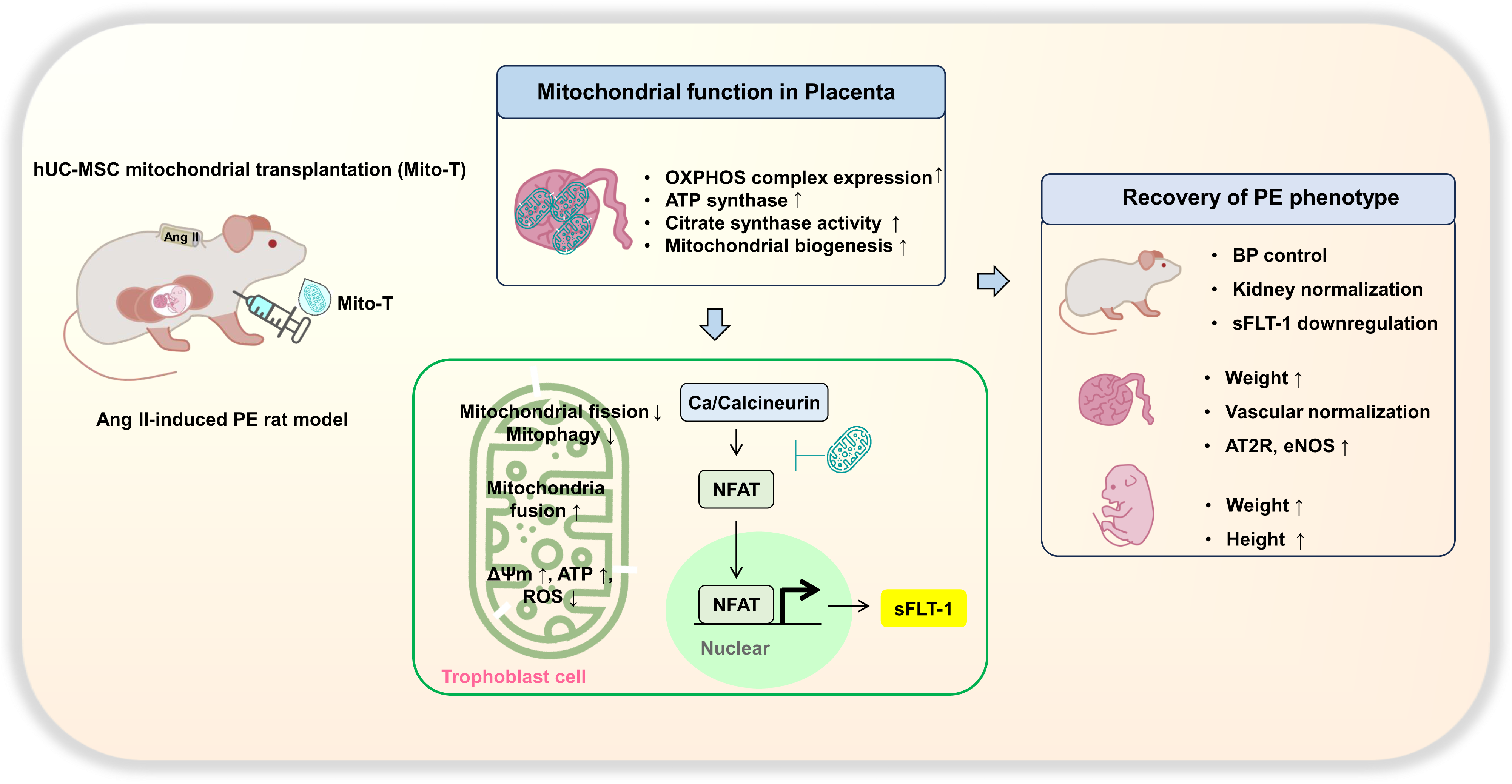
Graphical abstract of human umbilical cord mesenchymal stem cell-derived mitochondria improving maternal phenotype by improving placental mitochondrial function in an angiotensin II-induced rat preeclampsia model.

